# Improved vector toolkit for genome writing in mammalian cells

**DOI:** 10.64898/2026.03.15.711894

**Authors:** Kelly Barriball, Brianna Berrios, Camilla Coelho, Sudarshan Pinglay, Yu Zhao, Noor Chalhoub, Tiffany Tsou, John T. Atwater, Jef D. Boeke, Weimin Zhang, Ran Brosh

## Abstract

Efficient genome writing in mammalian cells requires robust methods for integrating large DNA payloads. The previously described method mammalian Switching Antibiotic resistance markers Progressively for Integration (mSwAP-In) enables iterative, biallelic genome rewriting in mammalian stem cells with DNA payloads exceeding 100 kb. However, the lack of standardized vectors and certain technical constraints have limited its broader adoption. Here we present an improved plasmid toolkit designed to streamline the implementation of mSwAP-In. The toolkit includes two core vectors. pLP-TK (pCTC174) is a landing-pad plasmid compatible with Golden Gate assembly of genomic homology arms and supports both mSwAP-In and the recombinase-mediated cassette exchange method Big-IN. mSwAP-In MC2v2 (pKBA135) is a versatile Big DNA assembly and delivery vector that supports Gibson-based assembly and incorporates positive, negative, and fluorescent selection markers, as well as a backbone counterselection cassette to minimize unwanted plasmid integration. The vector architecture also enables propagation in yeast and bacterial hosts, inducible plasmid copy-number amplification in standard E. coli strains, and CRISPR/Cas9-mediated payload release through preinstalled guide RNA target sites. We further characterize the FCU1/5-FC counterselection system in mouse embryonic stem cells and define conditions that minimize its bystander toxicity. Finally, we provide a set of Cas9-gRNA expression plasmids optimized for common mSwAP-In applications. Together, these reagents constitute a standardized and experimentally validated toolkit that simplifies large-scale genome writing using mSwAP-In.

## INTRODUCTION

Mammalian genome writing—the insertion or replacement of large (>10 kb) genomic sequences—is an emerging technology with the potential to revolutionize disease-associated haplotype dissection, animal model engineering, cell therapies, and biomanufacturing, among other fields^1^. We recently developed two complementary genome writing methods, Big-IN^2^ and mSwAP-In^3^, that enable the writing of genomic sequences at the ∼200 kb range in pluripotent stem cells. These methods have been used for a variety of applications, including the dissection of regulatory mechanisms of several key mouse developmental genes^4-7^ and a human Type 2 diabetes risk haplotype^8^, the study of default genomic states^9^, and the generation of authentic next-gen humanized mouse models for COVID-19^3^ and X-linked Dystonia Parkinsonism^10,11^.

Of particular relevance here is ‘mammalian switching antibiotic resistance markers progressively for integration’ (mSwAP-In). This method employs CRISPR/Cas9 to introduce double-stranded breaks in the genome, followed by homology-directed repair to mediate insertion of new DNA payloads, and a powerful combination of positive and negative selection markers (contained within “marker cassettes”) to isolate clonal populations with correct integrations^3^. By reusing a pair of marker cassettes (MC1 and MC2), mSwAP-In theoretically allows unlimited iterative genome writing. For example, in a particular delivery step, a DNA payload containing MC2 overwrites MC1 (integrated in the previous step). A positive selection marker (*e*.*g*., blasticidin resistance gene, BSD) is used to select for cells that integrated the incoming payload containing MC2, whereas a negative selection marker (*e*.*g*., thymidine kinase, TK) is used to select against cells that still retain MC1. In the next step, a second payload, this time containing MC1, can be used to overwrite MC2, as long as each cassette contains unique selection markers.

Despite its elegance and utility, a few technical details impede the wide adoption of mSwAP-In. These include: (1) the lack of a standard landing pad vector that harbors marker cassette MC1 and can be easily modified with locus-specific homology arms; (2) the lack of a standard payload acceptor vector that harbors marker cassette MC2 and can be easily modified with locus-specific homology arms and payload-specific assembly arms; (3) the use of the *HPRT1* gene as the MC2 negative (counterselectable) marker in the original iteration of mSwAP-In, which necessitates an additional genome engineering step (deleting the endogenous *HPRT1* gene) and in some cases, limits the generation of mouse models due to the neurological defects associated with HPRT inactivation^12^; (4) the requirement to use EPI300, an expensive commercial strain of *E. coli*, which enables plasmid copy number induction, for obtaining high concentration plasmid preps; and (5) the lack of counterselectable plasmid backbone markers, which can be deployed to minimize backbone integration and retention of episomal plasmids.

To overcome these limitations, we designed and tested two new plasmid vectors with enhanced properties. The first vector, pLP-TK (pCTC174), is a “landing pad” plasmid that carries MC1, whereas mSwAP-In MC2v2 (pKBA135) is a Big DNA assembly and delivery vector that carries an updated version of MC2, MC2v2. Both vectors have been thoroughly tested in our lab in terms of efficiency and ease of DNA assembly and delivery to mammalian cells. Combined, these two plasmids enable the facile deployment of genome writing using mSwAP-In.

## RESULTS AND DISCUSSION

### pLP-TK (pCTC174), a user-friendly landing pad vector compatible with mSwAP-In and Big-IN that harbors a counterselectable backbone

pLP-TK (pCTC174) is a landing pad plasmid designed for integration into mammalian genomes as an intermediate step prior to the delivery of Big DNA payloads. **Figure 1** depicts a SnapGene-generated map of the plasmid. Once integrated into the genome, the landing pad (LP) region of pLP-TK is compatible with both mSwAP-In and Big-IN due to the presence of terminal guide RNA (gRNA) binding sites that enable CRISPR/Cas9-mediated removal, and a pair of heterotypic loxM/loxP sites that enable recombinase-mediated cassette exchange (RMCE), respectively. pLP-TK carries a marker cassette MC1 which is compatible with the set of selection markers in both Big-IN and mSwAP-In payload vectors.

**Figure 1.**
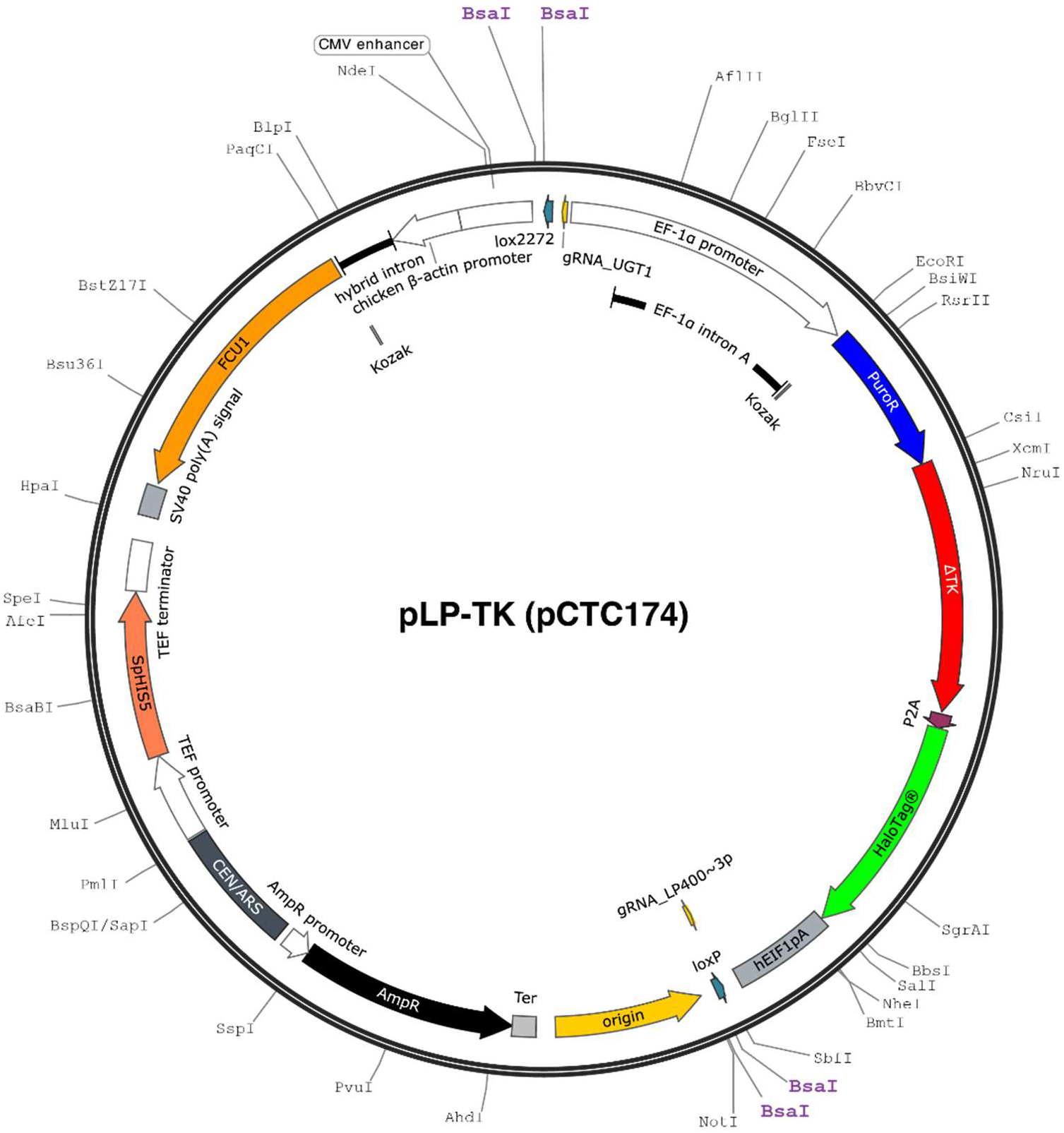
pLP-TK (pCTC174). Plasmid map, adapted from SnapGene. Unique RE sites are displayed on the outside in black. Four BsaI sites are indicated in purple. Components and elements starting from position 0 (clockwise): lox2272, Cre recombination site (also known as loxM); gRNA_UGT1, guide RNA binding site (see **Table 2**); EF-1α promoter, human *EEF1A1* gene promoter and 5’ UTR that includes an internal intron and Kozak sequence^30^; PuroR, *S. alboniger* Puromycin N-acetyltransferase; ΔTK, truncated herpes simplex virus thymidine kinase^14^; P2A, 2A peptide from porcine teschovirus-1 polyprotein^15^; HaloTag®, modified bacterial dehalogenase fluorescence marker^16^; hEIF1pA, human *EIF1* polyadenylation signal; gRNA_LP400∼3p, guide RNA binding site (**Table 2**); loxP, Cre recombination site (heterotypic to loxM); Ori, high-copy-number ColE1/pMB1/pBR322/pUC origin of replication; AmpR promoter—AmpR—Ter, Beta-lactamase bacterial expression cassette; CEN/ARS, *S. cerevisiae CEN6* centromere fused to an autonomously replicating sequence; TEF promoter—*SpHIS5*—TEF terminator, *Schizosaccharomyces pombe* imidazoleglycerol-phosphate dehydratase expression cassette, required for histidine biosynthesis and complements *S. cerevisiae his3* mutations; CMV enhancer—chicken β-actin promoter—hybrid intron—*FCU1*—SV40 poly(A), mammalian *FCU1* expression cassette. The combination of the human cytomegalovirus (CMV) immediate early enhancer, chicken β-actin promoter, and hybrid intron (hybrid between chicken β-actin and minute virus of mice^17^) constitutes the CAG promoter. The *FCU1* gene is an engineered, bifunctional fusion gene derived from yeast *FCY1* and *FUR1*^18^. SV40 poly(A) is the polyadenylation signal from SV40 virus.

Using a single Golden Gate assembly (GGA)^13^ reaction (**Fig. S1**), two homology arms (HAs) for mammalian genome targeting are cloned into the two pairs of mutually incompatible BsaI restriction enzyme (RE) sites. **Table 1a** lists primer templates for amplification of HAs compatible with pLP-TK’s BsaI GGA cloning strategy. To enhance LP on-target integration and limit backbone integration, gRNA target sites can be cloned externally to each HA in the same reaction^2^. Using the same gRNA target sites used to induce genomic cuts is recommended to minimize the number of gRNAs needed for each integration. **Table 1b** lists primer templates for amplifying HAs + gRNAs. Note that internal BsaI sites in the HAs can significantly reduce cloning efficiency, and it is recommended to avoid them if possible.

**Table 1a.** Landing pad homology arms cloning primer template. N(4), 4-nt 5’ clamp to ensure efficient BsaI digestion (can be shortened); Red, BsaI binding sites; Blue, BsaI cutting sites; N(20), ∼20-nt template-binding sequence (to amplify genomic region corresponding to the HA).

| Primer purpose | Primer sequence (5' – 3') |
| --- | --- |
| Amplification of HAL, Forward | N(4)-GGTCTCACCCT-N(20) |
| Amplification of HAL, Reverse | N(4)-GGTCTCACGTT-N(20) |
| Amplification of HAR, Forward | N(4)-GGTCTCGTATG-N(20) |
| Amplification of HAR, Reverse | N(4)-GGTCTCAGGAT-N(20) |

**Table 1b.** Landing pad homology arms and gRNA binding sites cloning primer template. For cloning HAs that are flanked with gRNA binding sites, use the template below. The 5’-gRNA and 3’-gRNA, as well as the PAM sequences, should all be cloned 5’ to 3’ (“facing right”). See Table 1a legend for details.

| Primer purpose | Primer sequence (5' – 3') |
| --- | --- |
| Amplification of HAL, Forward | N(4)-GGTCTCACCCT-N(20)-5'-gRNA-PAM-N20 |
| Amplification of HAL, Reverse | N(4)-GGTCTCACGTT-N(20) |
| Amplification of HAR, Forward | N(4)-GGTCTCGTATG-N(20) |
| Amplification of HAR, Reverse | N(4)-GGTCTCAGGAT-N(20)-3'-gRNA-PAM-N20 |

**Table 2.**
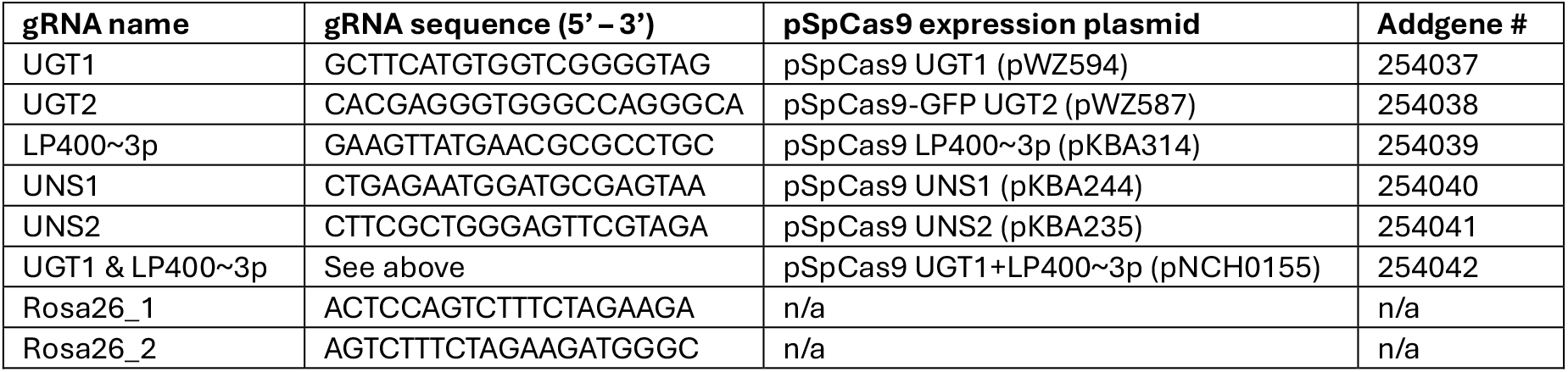
Guide RNAs (gRNAs) and expression vectors.

The genomically-integrated portion of pLP-TK (the landing pad, LP-TK) contains the following elements:

1. A terminal pair of heterotypic lox sites, lox2272 (aka loxM) on the left, and loxP on the right. These sites are compatible with the payload vectors used in Big-IN^2^, allowing Cre-mediated RMCE.
2. A terminal pair of gRNA sites, gRNA_UGT1 (universal gRNA target 1) and gRNA_LP400∼3p (see **Table 2** for gRNA sequences). Both sites are sequence-unique relative to the human and mouse genomes. This pair of gRNA targets is used to release the LP during mSwAP-In payload delivery. gRNA_UGT1 is also compatible with the design of mSwAP-In MC2v2 (pKBA135), as described below.
3. A single open reading frame (ORF) driven by a human EF-1α promoter and terminated by a human *EIF1* polyadenylation site. The ORF consists of a positive selection marker PuroR (*S. alboniger* Puromycin N-acetyltransferase) that renders cells resistant to puromycin, fused to a negative (counterselectable) truncated herpes simplex virus thymidine kinase (ΔTK) marker gene^14^, which renders cells sensitive to ganciclovir (GCV). A porcine teschovirus-1 P2A peptide^15^ separates the PuroRΔTK fusion protein from a HaloTag® marker^16^, which enables fluorescence-based detection or sorting of LP-TK-harboring cells.

pLP-TK’s plasmid backbone contains the following elements:

1. A high-copy-number ColE1/pMB1/pBR322/pUC origin of replication.
2. An ampicillin resistance marker cassette encoding Beta-lactamase.
3. An *S. cerevisiae* CEN6 centromere fused to an autonomously replicating sequence (ARS), as well as a *Schizosaccharomyces pombe his5* expression cassette. Combined, these elements enable propagation and selection in yeast host cells.
4. A CAG promoter^17^-driven counterselectable *FCU1* expression cassette. *FCU1* is a fusion of two *S. cerevisiae* genes, cytosine deaminase (*FCY1*) and uracil phosphoribosyltransferase genes (*FUR1*). When expressed, *FCU1* functions as a “suicide” gene that converts nontoxic 5-fluorocytosine (5-FC) into the toxic metabolites 5-fluorouracil (5-FU) and 5-fluorouridine-5’-monophosphate (5-FUMP)^18^.

The presence of the *FCU1* backbone marker cassette enables 5-FC-induced selective killing of cells with undesirable pLP-TK events, such as off -target integration, tandem on-target integration, and episomal propagation, all of which are usually accompanied by the retention of the plasmid backbone. However, treatment of pLP-TK-transfected cells with 5-FC can only be applied once the initial population of transfected plasmids has been turned over, which typically takes 5 or more days in mouse embryonic stem cells. Only at this point, clones in which the landing pad has been genomically integrated at a single copy and the backbone has been degraded would be 5-FC resistant. In addition, since the toxic derivatives of 5-FC can poison neighboring cells via the “bystander effect” (see below), it is recommended that the main population of engineered cells to be stored or used as the stock for future work is not treated with 5-FC. Instead, following clone picking, each clone is split into two duplicate plates, and only one is treated with 5-FC to identify clones in which the plasmid backbone is retained genomically or episomally. Such duplicate plating is already part of the standard engineering pipeline used when applying mSwAP-In for generating genomically engineered mouse models^3^.

pLP-TK (pCTC174) was deposited on Addgene as plasmid #254034

### mSwAP-In MC2v2 (pKBA135), a universal mSwAP-In payload assembly vector

mSwAP-In MC2v2 (pKBA135) (**Fig. 2**) is a multipurpose vector designed to support low-copy assembly and editing of Big DNA payloads in yeast or bacterial host cells, propagation and copy number induction in bacterial cells prior to purification, delivery to mammalian cells, and isolation of clones containing a correctly integrated payload. The vector carries marker cassette MC2v2, which is complementary to pLP-TK’s MC1.

**Figure 2.**
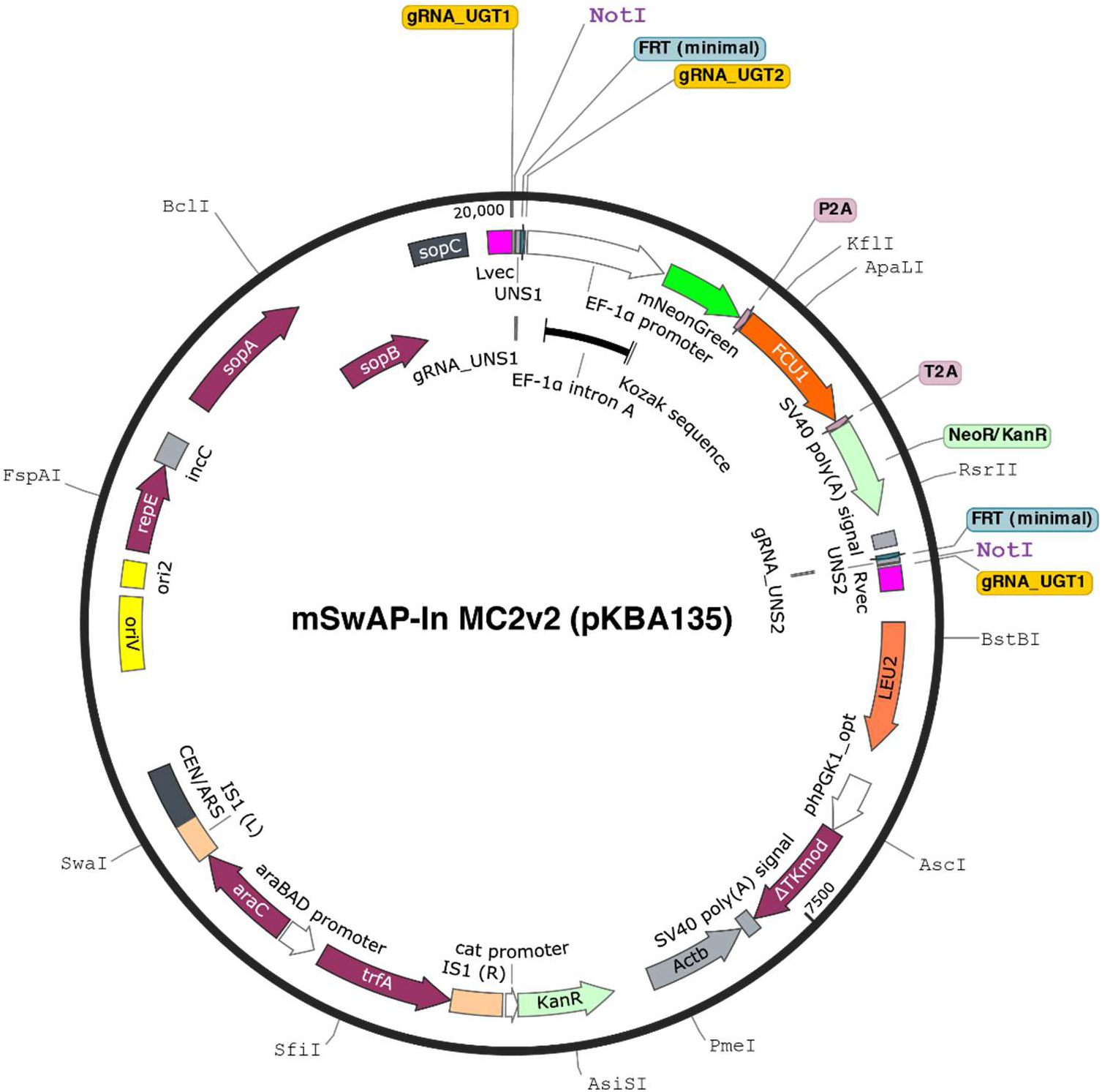
mSwAP-In MC2v2 (pKBA135) Plasmid map, adapted from SnapGene. Unique RE sites are displayed on the outside in black. Two NotI sites are indicated in purple. Components and elements starting from position 0 (clockwise): gRNA_UGT1, guide RNA binding site (**Table 2** for gRNA sequences); UNS1, unique nucleotide sequence 1 (Ref^31^); gRNA_UNS1, guide RNA binding site; FRT (minimal), FLP recombinase from the *S. cerevisiae* 2μ plasmid; gRNA_UGT2, guide RNA binding site, EF-1α promoter, human *EEF1A1* gene promoter and 5’ UTR that includes an internal intron and a Kozak sequence^30^; mNeonGreen, bright monomeric yellow-green fluorescent protein^32^; P2A, 2A peptide from porcine teschovirus-1 polyprotein^15^; *FCU1*, an engineered, bifunctional fusion gene derived from yeast FCY1 and FUR1^18^; T2A, 2A peptide from *Thosea asigna* virus; NeoR/KanR, neomycin (G418) resistance gene aminoglycoside phosphotransferase from Tn5; SV40 poly(A), SV40 virus polyadenylation signal; FRT (minimal), FLP recombinase from the *S. cerevisiae* 2μ plasmid; gRNA_UNS2, guide RNA binding site; UNS2, unique nucleotide sequence 2 (Ref^31^); gRNA_UGT1, guide RNA binding site; Rvec, universal linker for yeast assembly; *LEU2*, yeast selection marker 3-isopropylmalate dehydrogenase, required for leucine biosynthesis; phPGK1_opt—*ΔTKmod*—SV40 poly(A); mammalian expression cassette for truncated herpes simplex virus thymidine kinase (ΔTK)^14^. phPGK1 is derived from the human phosphoglycerate kinase 1 promoter and has been mutated to eliminate several RE sites. ΔTKmod has been synonymously recoded to eliminate RE sites; Actb, a region of the mouse *Actb* gene, nonfunctional, used for payload copy number quantification after delivery^3^; cat promoter—KanR, a kanamycin resistance bacterial expression cassette encoding aminoglycoside phosphotransferase; IS1(R), a truncated portion of the bacterial Insertion Sequence 1 (Ref^33^), araC— araBAD promoter—trfA, a bacterial arabinose-dependent copy number induction cassette^34^; IS1(L), a truncated portion of the bacterial Insertion Sequence 1; CEN/ARS, *S. cerevisiae CEN6* centromere fused to an autonomously replicating sequence; oriV, origin of replication for the bacterial F plasmid; ori2, secondary origin of replication for the bacterial F plasmid; also known as oriS; repE, replication initiation protein for the bacterial F plasmid; incC, incompatibility region of the bacterial F plasmid; sopA, partitioning protein for the bacterial F plasmid; sopB, partitioning protein for the bacterial F plasmid; sopC, centromere-like partitioning region of the bacterial F plasmid; Lvec, universal linker for yeast assembly.

A two-step assembly strategy (**Fig. 3**) is often used to generate a delivery-ready vector harboring a Big DNA payload. First, pKBA135 is digested into two fragments using a NotI RE. Then, a Gibson assembly^19^ reaction is used to reassemble the vector with a pair of homology arms for mammalian genomic integration. In the same step, an “assembly fragment” is added. The assembly fragment comprises two short (∼200 bp) assembly arms, separated by a unique RE site (typically SalI). In the second step, the acceptor vector is linearized using SalI (or any other unique RE that cuts between the assembly arms), purified, and used for assembly of the payload. The payload assembly can be done in at least three separate ways. First, lambda Red recombination^20^ in an *E. coli* strain that carries a bacterial artificial chromosome (BAC) can be used to quickly clone any region of the BAC. Second, a BAC can be purified from *E. coli* and transformed along with the digested acceptor vector into yeast (*S. cerevisiae*) host cells. A third option is using the digested acceptor vector for multi-segment assembly in yeast. In all three, the assembly arms AAL and AAR within the assembly fragment dictate the left and right boundaries of the payload. Backbone selection markers enable isolation of circular plasmids following assembly in bacteria (*KanR*) or yeast (*LEU2*).

**Figure 3.**
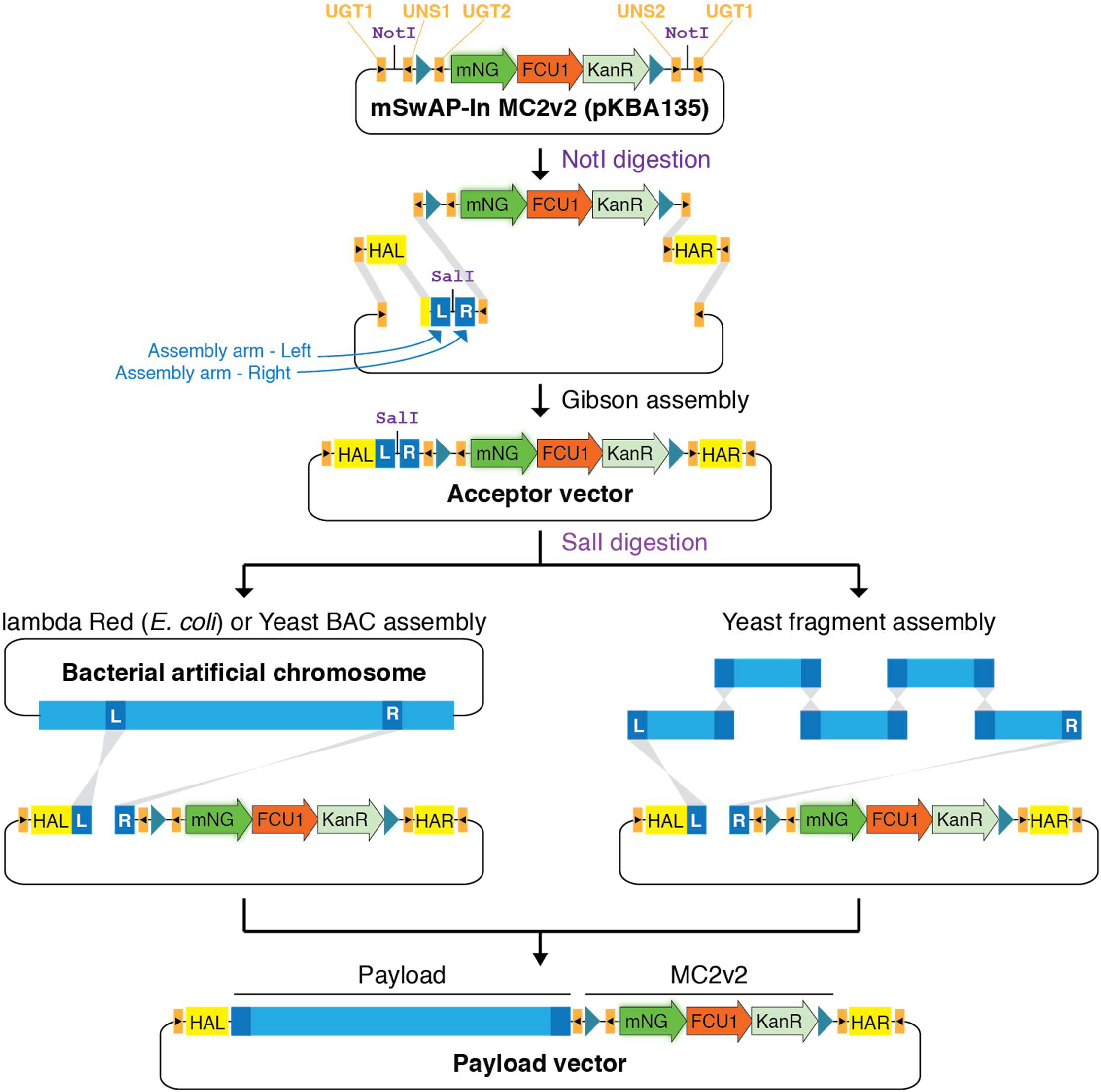
Payload assembly into mSwAP-In MC2v2 (pKBA135) Top, simplified schematic of MC2v2 (pKBA135) using the same color scheme as in Figure 2. Backbone elements are not included. MC2 elements include gRNA binding sites (orange boxes with orientation indicated by a black arrow), FRT sites (blue triangles), mNeonGreen (mNG), FCU1 and KanR. Middle, assembly of a delivery-ready Big DNA payload into pKBA135 is performed in two steps. First, pKBA135 is digested with NotI and Gibson assembly is used to install two homology arms (HAL and HAR) for genome integration and an “assembly fragment” that contains two assembly arms – assembly arm left, AAL (blue box labeled “L”), and assembly arm right, AAR (blue box labeled “R”). AAL and AAR are separated by a unique restriction enzyme; in this case, SalI. Each fragment includes short terminal homologies to its neighboring fragments to enable Gibson assembly. Note that the assembly fragment can also be cloned on the right side of MC2, which would reverse the order of the payload relative to MC2. In the second step, the resulting acceptor vector is linearized with SalI (or any other unique cutter preinstalled in the assembly fragment) and assembly with the payload is performed. Payload DNA can be sourced as a single fragment from bacterial artificial chromosomes (BACs) and assembled with the linearized acceptor vector in yeast or bacterial hosts. Alternatively, multi-fragment yeast assembly can be performed using PCR products or synthetic DNA fragments that include short terminal homologies to neighboring fragments (dark blue boxes). Bottom, fully assembled delivery-ready payload vector. Plasmid backbone elements are excluded for simplicity.

Similar to standard BACs and the previous generation of mSwAP-In payload vectors, pKBA135 harbors the backbone elements that are required for stable, low-copy propagation in bacterial host cells, including the *E. coli* F plasmid-derived sopA, sopB, sopC, incC and ori2^21^. Additionally, the backbone of pKBA135 harbors a copy number induction (CNI) cassette, which includes the trans-acting replication protein *trfA*, L-arabinose regulatory protein *araC*, and the promoter of the L-arabinose operon of *E. coli* (*araBAD* promoter). Treatment with L-arabinose induces the expression of *trfA*, which binds to and activates oriV, leading to increased replication. Thus, the CNI cassette enables arabinose-dependent amplification of pKBA135 and payload-containing derivatives thereof. Unlike previous plasmid versions, this design makes CNI feasible in standard *E. coli* host strains such as Top10 and DH5α, avoiding dependence on expensive commercial strains such as EPI300^22^. The CNI cassette was cloned into the MluI site in an Insertion Sequence 1 (IS1) element that transposed into a previous parental version of the plasmid backbone, functionally splitting IS1 into two halves, IS1(L) and IS1(R). **Figure S2** illustrates CNI efficiency in a standard Top10 bacterial host strain.

To enhance the efficiency and fidelity of mammalian genome integration, pKBA135 was equipped with a backbone negative (counterselectable) marker cassette. The cassette consists of a human phosphoglycerate kinase 1 (*PGK1*) promoter driving the expression of a herpes simplex virus thymidine kinase (ΔTK)^14^ gene, followed by an SV40 polyadenylation signal. To eliminate several RE binding sites, both the promoter and the ΔTK gene were modified. Specifically, for the human *PGK1* promoter we used the mouse orthologous sequence to inform on nucleotide positions that are likely to tolerate change, while the ΔTK coding sequence was synonymously mutated. Following delivery to mammalian cells, selection with ganciclovir (GCV) leads to killing of cells that retain the ΔTK-harboring plasmid backbone, helping to reduce off -target and tandem on-target integration events, as well as episomal propagation of the payload plasmid.

Delivery fidelity is also aided by the presence of two UGT1 gRNA binding sites immediately flanking the payload and HAs (**Fig. 3**). This architecture is compatible with pLP-TK (pCTC174), which contains a UGT1 binding site in its left LP terminus. When overwriting a genomically integrated MC1, gRNA_UGT1 mediates one cut at MC1 and two cuts at the incoming payload vector, effectively releasing the payload from its backbone. A second gRNA, LP400∼3p, cuts at the right terminus of LP-TK, facilitating its complete removal. **Table 2** lists relevant gRNA sequences.

Following delivery of an MC2v2-harboring payload vector, positive selection with G418 (Geneticin®) is used to isolate cells with NeoR-harboring MC2v2 genomic integration. Negative selection with GCV, which is typically applied starting day ∼5 post-transfection^3^, doubly selects against cells that express ΔTK either from MC1/LP-TK and/or from pKBA135’s plasmid backbone. Alternatively, or additionally, fluorescence-activated cell sorting/picking can be used to enrich for cells in which MC1 was overwritten by MC2v2. Such cells will lose HaloTag® expression and gain mNeonGreen.

MC2v2 contains a negative selection *FCU1* gene; its mechanism of function is described above. Selecting for the loss of MC2v2 is useful in two scenarios. First, following payload delivery, active transcription of the EF1α-driven marker cassette might interfere with downstream applications^23^, such as the dissection of regulatory mechanisms^1^. In this case, a subsequent “scar removal” step can be performed to excise MC2v2 from the genome. To this end, a pair of minimal FRT sites is situated flanking the cassette, allowing its excision by Flp recombinase. This approach would typically leave a 114-bp minimal scar in the genome, consisting of the UNS1, FRT and UNS2 elements; yet these sequences are not expressed and should have minimal effect on the engineered locus behavior. **Table 3** lists sequences of the aforementioned elements. An alternative scar removal approach is using CRISPR/Cas9 to initiate two genomic cuts at the termini of MC2v2 and providing a repair donor for complete removal of the scar. To this end, a pair of gRNA binding sites, gRNA_UNS1 and gRNA_UNS2, can be used. In both cases, successful scar removal can be enriched by the selective killing of cells that still express *FCU1* using 5-FC, with the bystander effect caveat described above. The second scenario where FCU1/5-FC negative selection against MC2v2 would be applied is the delivery of a second Big DNA payload using mSwAP-In’s iterative capability. In this case, the second payload would be designed to overwrite MC2v2 and introduce a version of MC1 to the genome.

**Table 3.**
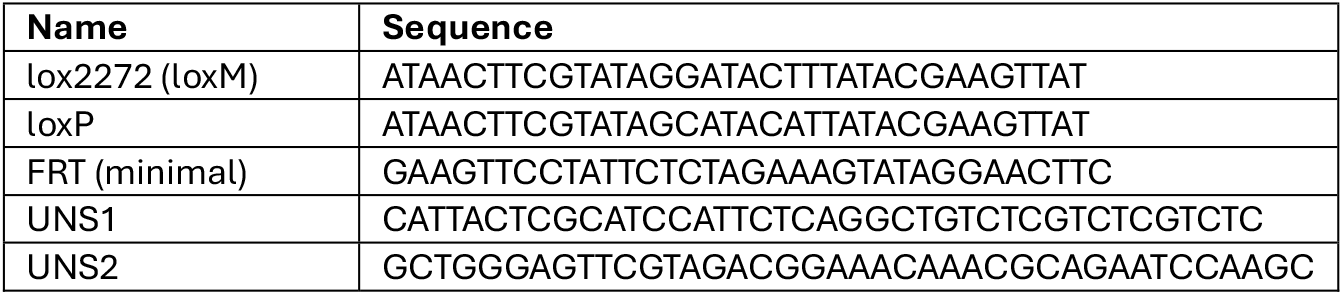
Sequences of selected plasmid elements.

In a previous study,^3^ we demonstrated efficient MC2 removal using the PiggyBac (PB) system^24^. While pKBA135 lacks the PB inverted terminal repeat (ITR) sequences, those short elements can be added during the cloning of the acceptor vector. PB-mediated excision of MC2 is completely scarless and does not require a donor repair template.

mSwAP-In MC2v2 (pKBA135) was deposited on Addgene as plasmid #254035

### Characterization of TK/GCV and FCU1/5-FC negative selection strategies

Application of mSwAP-In for iterative genome writing requires the use of two marker cassettes (MCs) with a distinct set of positive and negative selection markers^3^. Since both TK and FCU1 can metabolize 5-FU^25^, we wanted to ensure that their combined usage would not lead to cross-toxicity. Additionally, we wanted to compare their relative efficiency in producing positively edited cells via mSwAP-In. To test this, we engineered two isogenic mouse embryonic stem cell (mESC) lines in which a marker cassette harboring either TK (MC-TK) or FCU1 (MC-FCU1) was integrated at a single copy into the X chromosome. We performed a kill curve analysis for both GCV and 5-FC for these two mESC lines and confirmed that each line is only sensitive to the drug corresponding to its negative selection marker, *i*.*e*., that MC-TK cells are exclusively sensitive to GCV while MC-FCU1 are exclusively sensitive to 5-FC (**Fig. 4a**). Next, to test the compatibility of FCU1/5-FC negative selection with the mSwAP-In schema, we delivered an empty payload plasmid harboring a NeoR gene and an eGFP gene to both cell lines and compared survival following application of positive selection only (Pos), negative selection only (Neg), and a combination of positive and negative selection (Pos+Neg). G418 was used for positive selection for both cell lines, and GCV and 5-FC were used for negative selection for MC-TK and MC-FCU1, respectively. We performed this experiment under two cell density conditions. After 9 days, cells were analyzed on a flow cytometer for viability (**Fig. 4b,c**). Interestingly, while viability following positive selection was similar for all 4 conditions, negative selection led to a marked decrease in viability of 5-FC-treated MC-FCU1 cells when plated at high density. This bystander effect was mostly avoided in the Pos+Neg condition since even for the dense condition, the initial positive selection led to a reduction in density by the time negative selection was applied.

**Figure 4.**
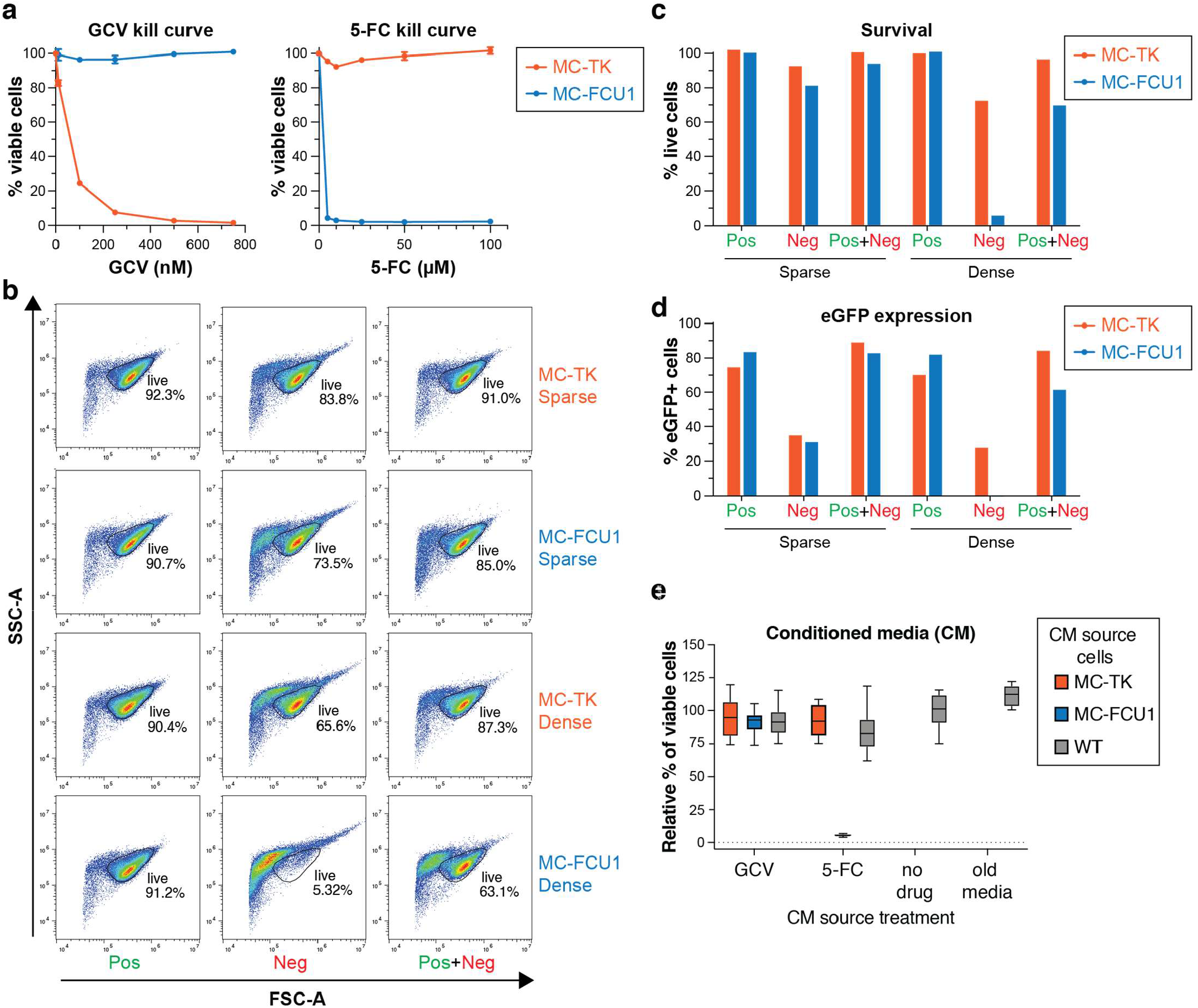
Characterization of negative selection with TK/GCV and FCU1/5-FC. **a**. mESCs engineered with a marker cassette expressing TK (MC-TK) or FCU1 (MC-FCU1) were seeded in 24-well plates at 4×10^4^ cells/well. One day later, cells were treated with the indicated concentrations of GCV or 5-FC for 3 days. Viability was measured using PrestoBlue. Experiment was performed as biological duplicates, measured as technical triplicates, and data are represented as mean +/-standard deviation. **b-c**. 3×10^6^ MC-TK or MC-FCU1 mESCs were nucleofected and seeded into two 10-cm plates at 90% and 10% of the total cell number, generating dense (2.7×10^6^ cells/plate) and sparse (3×10^5^ cells/plate) plating conditions. Three different selection schemes were compared: Positive selection only (Pos) included treatment with 150 µm/mL G418 starting day 1 post-seeding; negative selection only (Neg) included treatment with either 250 nM GCV or 5µM 5-FC, for MC-TK and MC-FCU1, respectively. Selection was applied starting day 3 post-seeding. Combinatorial positive and negative selection (Pos+Neg) included 150 µm/mL G418 starting day 1 post-seeding and GCV/5-FC at 250 nM/5µM starting day 3 post-seeding. Analysis was performed on day 9 post-seeding by analyzing cells on a flow cytometer for forward scatter (FSC-A) and side scatter (SSC-A) and gating on live cells (panel b, quantified in panel c). **d**. Live cells (panel b) were analyzed for eGFP expression. **e**. Effect of conditioned media on cell viability. Confluent cultures of MC-TK, MC-FCU1 and parental (WT) cells were treated with the indicated drugs for 30 hours. Conditioned media (CM) was collected, filtered, diluted 1:1 with fresh media and used to grow WT cells for a total of 3 days. Viability was measured with PrestoBlue. Experiment was performed in biological triplicates, measured in technical triplicates, and data are represented as mean +/-standard deviation after normalizing to the “no drug” condition. Old media served as a control for the effect of spent media.

Ultimately, what matters most is the number of correctly edited clones. When comparing the fraction of eGFP+ clones (**Fig. 4d**), which is indicative of payload plasmid presence, we note that under dense plating conditions, 5-FC selection performs poorly. However, under sparse plating conditions, we see a comparable percentage of eGFP+ clones for GCV and 5-FC selection, indicating that the toxic bystander effect is mostly mitigated by a lower plating density. Importantly, both negative selection strategies performed very well following positive selection, especially under sparse conditions, where over 80% of the cells were eGFP+.

Finally, to better understand the mechanism of the observed FCU1/5-FC bystander effect, we performed an experiment to test whether this toxicity can be mediated by diffusion of toxic metabolites. Specifically, we treated dense cultures of MC-TK, MC-FCU1 and parental wild type (WT) cells with GCV or 5-FC for 30 hours. We then collected this conditioned media (CM), filtered it, mixed it 1:1 with fresh media, and used it with WT cells for a total of 3 days, after which we analyzed their viability. As shown in **Figure 4e**, WT cells exposed to media conditioned on 5-FC-treated MC-FCU1 cells died completely, while all other conditions, including CM from GCV-treated MC-TK cells, had no significant effect on viability. This result clearly demonstrates that, unlike TK/GCV, the FCU1/5-FC bystander effect does not require cell-cell contacts and clarifies why it is crucial to maintain low density when applying it. Notably, even at low density, it is possible that FCU-negative cells might acquire DNA damage when exposed to toxic derivatives of 5-FC, which is why we recommend duplicate plating as described above.

### pKBA204, a Rosa26-targeting MC2v2 acceptor vector

The Rosa26 safe harbor locus is one of the most popular choices for integrating DNA payloads into the mouse genome^26,27^. We generated mSwAP-In_MC2v2 (Rosa26)_Acceptor (pKBA204), a derivative version of mSwAP-In MC2v2 (pKBA135) with Rosa26-targeting homology arms. Specifically, a 1.6-kb HAL corresponds to mm10 chr6:113074387-113076025, while a 1.2-kb HAR corresponds to chr6:113076031-113077230. pKBA204 contains two unique RE sites, AvrII and PacI, located between HAL and MC2v2, which allow RE cloning of an acceptor fragment for downstream assembly of a Rosa26-targeting payload vector. pKBA204 is compatible with two pCas9 expression vectors that express Rosa26-targeting gRNAs, pSpCas9-BSD Rosa26_1 (pJA014) and pSpCas9 mRosa26_2 (pWZ869) (see **Table 2**).

mSwAP-In_MC2v2(Rosa26)_Acceptor (pKBA204) was deposited on Addgene as plasmid #254036

### A series of pSpCas9+gRNA expression vectors compatible with mSwAP-In applications

We cloned a series of expression vectors that are based on the architecture of pX330, a mammalian dual expression vector harboring a CAG promoter-driven SpCas9 gene and a U6 promoter-driven gRNA^28^. **Table 2** lists the gRNAs cloned into each vector and their corresponding Addgene plasmid numbers. These include gRNAs that target various parts of MC1 and MC2v2 (UGT1, UGT2, LP400∼3p, UNS1, and UNS2) as well as two gRNAs that target the mouse Rosa26 locus and are compatible with the aforementioned pKBA204.

We also cloned pNCH0155, a vector that expresses SpCas9 and both gRNA_UGT1 and gRNA_LP400∼3p. To this end, we have used pX333 (Addgene plasmid # 64073) a vector for tandem expression of two sgRNAs from two independent U6 promoters^29^.

## METHODS

### mESC culturing

Both mESC lines were cultured on plates coated with 0.1% gelatin in ESM +2i media. Composition of ESM +2i media as follows: KnockOut DMEM supplemented with 15% FBS, 0.1 mM 2-mercaptoethanol, 1% GlutaMAX, 1% MEM non-essential amino acids, 1% penicillin-streptomycin, 1,250 U ml^−1^ LIF, 3 μM CHIR99021, and 1 μM PD0325901. Media was changed daily, and cells were passaged at approximately 85% confluency with Accutase, incubated at 37 °C for 8 minutes.

### mESC nucleofection

Cells were seeded at 2×10^6^ cells/10 cm plate two days before nucleofection. On day 0, plates were dissociated using 5 mL Accutase, and the resulting cell count was determined using the ThermoFisher Countess 3. 3×10^6^ were nucleofected with 3 µg empty payload plasmid and 1.5 µg of each pSpCas9 plasmid. 6 mL ESM +2i media was used to recover cells and then each nucleofection was split into 3 selection conditions (+, -, ±), 2mL each, and each condition was further split into 10/90% sparse/dense plating. All 6 conditions for each cell line were plated on 10 cm pre-gelatinized dishes (12 plates total). Plates were given 8 mL ESM +2i media total and left incubating at 37 °C overnight. Positive selection was with 150 µg/mL G418 starting 24 hours after nucleofection. Negative selection was with 250 nM GCV for MC-TK cells and 5 µM 5-FC for MC-FCU1; both were started 72 hours after nucleofection.

### PrestoBlue viability assay

Cell plates were rinsed with PBS and then incubated with 1X PrestoBlue (ThermoFisher Scientific A13262, diluted 1:10 in DMEM) at 37 °C for 20 minutes. For 24-well and 48-well plates, 250 µL/well and 500 µL/well PrestoBlue were used, respectively. PrestoBlue solution was transferred to a black-walled, clear-bottomed 96-well plate as technical triplicates and measured using a Synergy H1 microplate reader at 560/590 excitation/emission profile as recommended.

### Flow analysis

Cells were dissociated with Accutase and resuspended in 1X PBS with 5% fetal bovine serum (FBS) before transferring to 40 µm strainer top flow cytometry tubes. Cell cultures were analyzed on a Sony Cell Analyzer with the following parameters: all 4 lasers on, sample flow rate 1.0, Threshold CH FC, Threshold value 25%, FSC Gain 10, SSC Voltage 30%, Fluorescence PMT voltage 50%. For quantifying eGFP+ cells, a negative control sample was used to establish a baseline threshold. Analysis was performed using the FlowJo software.

### Conditioned media

Each mESC line (MC-TK, MC-FCU1, and parental (WT)) was plated at high confluency (2.2×10^6^ cells/plate) onto a pair of 10 cm TC-treated plates coated with 0.1% gelatin. Each plate received 9.45 mL of ESM +2i media with one of either drug added at the same concentration as previously: 250 nM GCV or 5 µM 5-FC. An additional WT plate received media with no drug. After 30 hours of media conditioning at 37 °C, media was harvested, filtered through a 0.22 µm filter and stored at 4 °C. WT cells were plated at a density of 3×10^3^ cells/well in a 48-well plate in ESM +2i media and allowed to adhere overnight at 37 °C. The next day, each well of 48-well plate of WT cells received 50% conditioned media, 50% fresh ESM +2i media, arranged as biological triplicates, and left to grow at 37 °C. After 72 hours, cell viability was measured following PrestoBlue viability assay as outlined above.

## AUTHOR CONTRIBUTIONS

C.C., S.P. and Y.Z. cloned pLP-TK (pCTC174). W.Z., B.B. and K.B. cloned mSwAP-In MC2v2 (pKBA135). W.Z. performed the copy number induction experiment. B.B. performed the TK/GCV and FCU1/5-FC characterization experiments. R.B. designed the pLP-TK (pCTC174) and mSwAP-In MC2v2 (pKBA135) vectors. W.Z. cloned pSpCas9 UGT1 (pWZ594) and pSpCas9-GFP UGT2 (pWZ587). K.B. cloned pSpCas9 LP400∼3p (pKBA314), pSpCas9 UNS1 (pKBA244) and pSpCas9 UNS2 (pKBA235). N.C. cloned pSpCas9 UGT1+LP400∼3p (pNCH0155). R.B., B.B. W.Z. and J.B. wrote the manuscript. N.C. and J.T.A. helped edit the manuscript.

## COMPETING INTERESTS

J.D.B and W.Z. are inventors on a pending patent application for mSwAP-In. J.D.B and R.B. are inventors on a pending patent application for Big-IN. J.D.B. is a Founder and Director of CDI Labs, Inc., a Founder of and consultant to Opentrons LabWorks/Neochromosome, Inc, a Founder of JATech, LLC, and serves or served on the Scientific Advisory Board of the following: CZ Biohub New York, LLC; Logomix, Inc.; Rome Therapeutics, Inc.; SeaHub, Seattle, WA; Tessera Therapeutics, Inc.; and the Wyss Institute.

## ACKNOWLEDGEMENTS

This work was supported in part by grant RM1HG009491 from the NHGRI Center for Excellence in Genome Science (CEGS), and grant 5R24OD037800 from the NIH Office of the Director, and by the Collaborative Center for X-Linked Dystonia-Parkinsonism (CCXDP).

## SUPPLEMENTAL FIGURES

**Figure S1.**
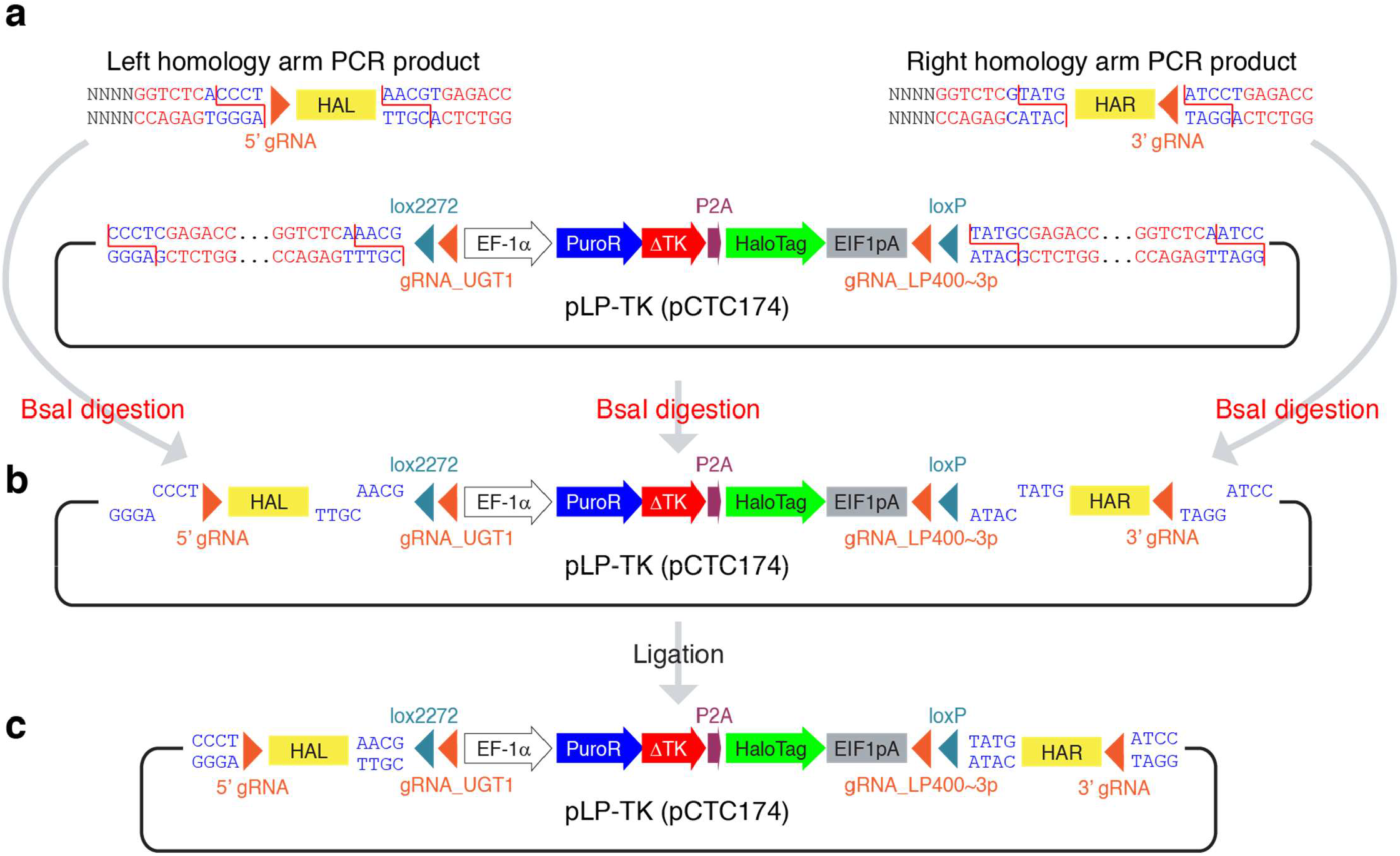
pLP-TK (pCTC174) Golden Gate assembly strategy. **a**. Schematic view of pLP-TK (pCT174) and PCR-amplified left homology arm (HAL) and right homology arm (HAR) with a focus on the sequences that comprise the BsaI restriction enzyme recognition sites (red) and the BsaI cut site (blue). A broken red line indicates the exact cutting pattern. The homology arms also include terminal gRNA cut sites for Cas9-mediated plasmid linearization. See **Tables 1a and 1b** for homology arm amplification primers. **b**. The result of BsaI digestion, which leaves 4-nt overhangs on each terminus of each sequence. **c**. A pLP-TK plasmid with homology arms for genomic integration is assembled following ligation, which is specified by the complementary 4-nt overhangs. Plasmid backbone elements are excluded for simplicity.

**Figure S2.**
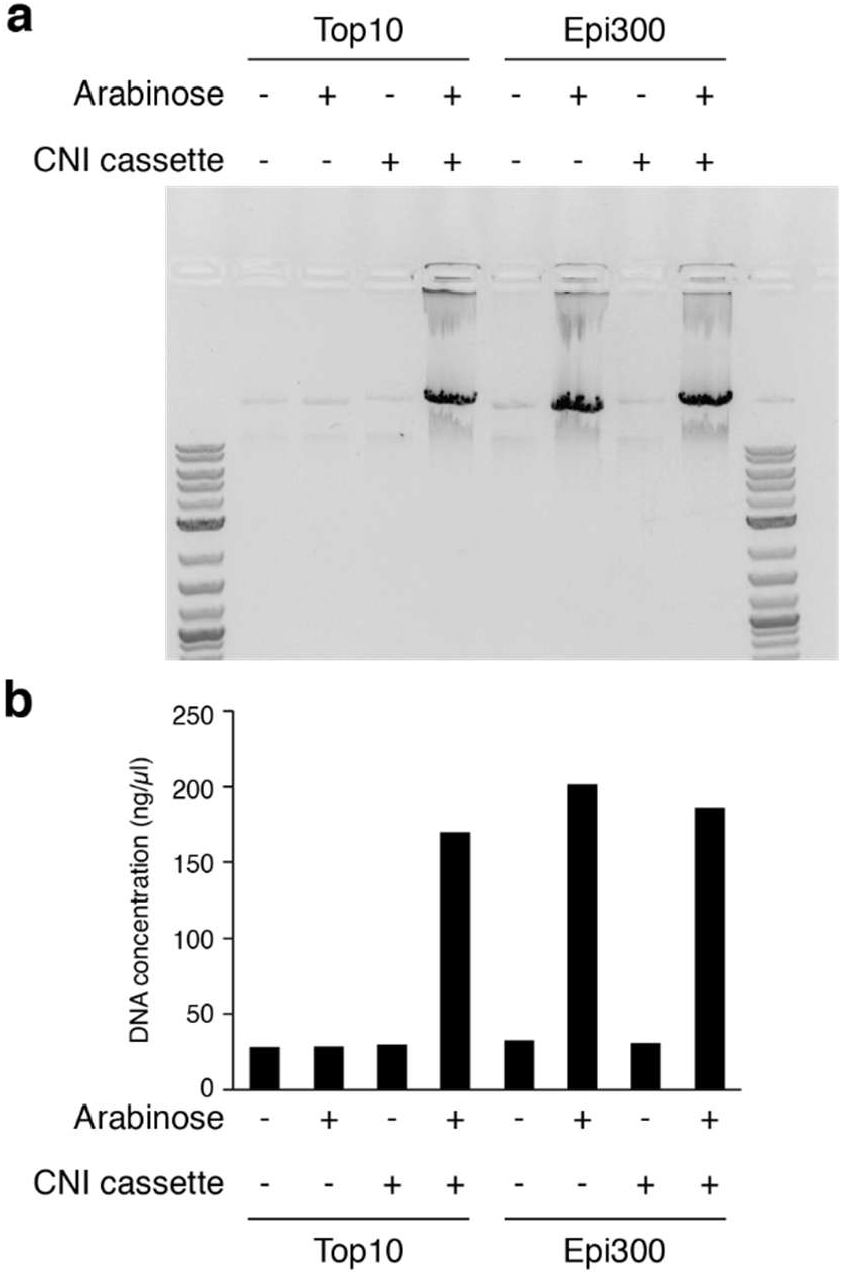
Arabinose-induced copy number induction. Plasmids lacking or containing a copy number induction (CNI) cassette (araC—araBAD promoter—trfA) were transformed into Top10 and EPI300 *E. coli* cells. EPI300 cells harbor a genomic CNI cassette^22^, whereas Top10 do not. After overnight growth with or without Arabinose, plasmids were miniprepped. **a**. Plasmid preparations were analyzed on a 1% Agarose gel and stained with EtBr. **b**. DNA concentration was measured using a NanoDrop instrument.

## Notes

### Summary of Updates

Author added (Camila Coelho), affiliations updated, and minor edits made

